# Treated HIV Infection Alters Phenotype But Not HIV-specific Function of Peripheral Blood Natural Killer Cells

**DOI:** 10.1101/2020.04.12.038604

**Authors:** Nancy Q. Zhao, Anne-Maud Ferreira, Philip M. Grant, Susan Holmes, Catherine A. Blish

## Abstract

Natural killer (NK) cells are the predominant antiviral cells of the innate immune system, and may play an important role in acquisition and disease progression of HIV. While untreated HIV infection is associated with distinct alterations in the peripheral blood NK cell repertoire, less is known about how NK phenotype is altered in the setting of long-term viral suppression with antiretroviral therapy (ART), as well as how NK memory can impact functional responses. As such, we sought to identify changes in NK cell phenotype and function using high-dimensional mass cytometry to simultaneously analyze both surface and functional marker expression of peripheral blood NK cells in a cohort of ART-suppressed, HIV+ patients and HIV-healthy controls. We found that the NK cell repertoire following IL-2 treatment was altered in individuals with treated HIV infection compared to healthy controls, with increased expression of markers including NKG2C and CD2, and decreased expression of CD244 and NKp30. Using co-culture assays with autologous, *in vitro* HIV-infected CD4 T cells, we identified a subset of NK cells with enhanced responsiveness to HIV-1-infected cells, but no differences in the magnitude of anti-HIV NK cell responses between the HIV+ and HIV-groups. In addition, by profiling of NK cell receptors on responding cells, we found similar phenotypes of HIV-responsive NK cell subsets in both groups. Lastly, we identified clusters of NK cells that are altered in individuals with treated HIV infection compared to healthy controls, but found that these clusters are distinct from those that respond to HIV *in vitro*. As such, we conclude that while chronic, treated HIV infection induces a reshaping of the IL-2-stimulated peripheral blood NK cell repertoire, it does so in a way that does not make the repertoire more HIV-specific.

## INTRODUCTION

Natural killer (NK) cells are critical effector cells of the innate immune system that can rapidly recognize and kill virally infected and tumour cells. NK cells express an array of activating and inhibitory receptors; the integration of signalling between these receptors determines NK cell activation and functional activity. This includes the release of cytolytic granules to induce target cell apoptosis, as well as the secretion of cytokines and chemokines including IFN-TNF-γ, MIP-1β (CCL4), and TNF-α.

Epidemiological and experimental evidence have highlighted the role of NK cells in the acquisition and disease progression of HIV-1. Increased constitutive NK cell activity is associated with protection from HIV acquisition in highly exposed seronegative individuals (Scott-Algara et al., 2003; Ravet et al., 2007). Similarly, the expression of the NK cell receptor KIR3DL1 and its cognate ligand HLA-Bw4-80I is associated with slower disease progression and improved suppression of autologous HIV-infected CD4 T cells (Martin et al., 2002; Alter et al., 2007; Song et al., 2014). Specific NK cell receptors have also been implicated in HIV recognition and targeting. NKG2A-expressing NK cells have improved activity against HIV (Lisovsky et al., 2015; Davis et al., 2016; Ramsuran et al., 2018), and NKG2D ligands are upregulated on HIV-infected cells (Richard et al., 2010). As such, changes in expression of NK cell receptors can impact their ability to target HIV.

Chronic, untreated HIV infection is associated with significant changes in the NK cell repertoire, the most well-defined of which is the loss of CD56^+^ NK cells, and the concomitant expansion of a CD56^neg^ NK cell subset (Alter et al., 2005). CD56^neg^ NK cells are functionally impaired and thought to be exhausted, demonstrating reduced cytotoxicity and IFN-TNF-γ production (Hu et al., 1995; Mavilio et al., 2005; Milush et al., 2013). In addition, the expression of the inhibitory receptor Siglec-7 (Brunetta et al., 2009), as well as the expression of the activating receptors NKp30, NKp44 and NKp46 (Mavilio et al., 2003), are decreased in chronic, viraemic HIV infection, whereas the expression of the inhibitory receptor TIGIT is increased (Yin et al., 2018; Vendrame et al., 2020). After treatment with antiretroviral therapy (ART), the patterns of CD56^+^ and CD56^neg^ NK cell subsets are restored to levels similar to seronegative, healthy individuals (Mavilio et al., 2005). However, less is known regarding how other NK cell subsets, as well as how the NK cell repertoire as a whole, may be altered in the setting of virological control by ART. In addition, the functional outcomes of these alterations, in particular with regards to how they may impact HIV-specific responses, are not well understood.

Contrary to their classic designation as an innate immune cell type, recent work has demonstrated the ability of human NK cells to form memory against viruses including cytomegalovirus, Epstein-Barr virus and varicella-zoster virus (Gumá et al., 2004, 2006; Lopez-Verges et al., 2011; Foley et al., 2012; Paust et al., 2017; Hammer et al., 2018; Nikzad et al., 2019). In non-human primates, infection with simian immunodeficiency virus (SIV) or SHIV generates antigen-specific NK cells that react with presented Gag and Env. In addition, vaccination with Ad26 vectors containing Gag and Env antigens from HIV and SIV generates long-lived, antigen-specific NK cells, even in the absence of continuous antigen stimulation (Reeves et al., 2015), raising the possibility that human NK cells in infected individuals could be similarly capable of generating and retaining memory responses against HIV antigens even without ongoing viral exposure. As such, we sought to understand whether previous HIV infection altered the functional capacity of peripheral blood NK cells to respond against a second, *in vitro* stimulation with autologous HIV-infected cells. Here, we use mass cytometry to profile NK cell receptor expression on a cohort of ART-suppressed, HIV+ donors and healthy controls, to determine how changes in the NK cell repertoire that occur with HIV infection influence HIV-specific NK cell responses.

## MATERIALS AND METHODS

### Study subjects and sample processing

Cryopreserved peripheral blood mononuclear cells (PBMCs) from HIV-infected patients treated with antiretroviral therapy (ART) were obtained from the Stanford HIV Aging Cohort. This study was approved by the Institutional Review Board of Stanford University. For anonymous healthy HIV uninfected donors, leukoreduction system chambers were obtained from the Stanford Blood Bank. PBMCs were isolated by density gradient centrifugation using Ficoll-Paque PLUS (GE Healthcare), and cryopreserved in 10% DMSO (Sigma Aldrich) and 90% fetal bovine serum (FBS, Thermo Fisher).

### CD4 and NK cell sorting and cell culture

PBMCs were thawed, and stained with a panel consisting of 7-AAD viability staining solution (eBioscience), CD14-BV421 (clone M5E2), CD19-BV421 (clone HIB19), CD16-FITC (clone 3G8), CD3-PE (clone SK7), CD4-BV711 (clone OKT4), and CD56-PE Cy7 (clone HCD56, all antibodies from Biolegend), and sorted for CD4 T cells (CD14^−^ CD19^−^ CD3^+^ CD4^+^) and NK cells (CD14^−^ CD19^−^ CD3^−^ CD56/CD16^+^) using a Sony SH800 sorter. Post-sorting, all cells were cultured in RPMI (Gibco), with 10% FBS (Thermo Fisher), 1% L-glutamine (Hyclone) and 1% penicillin/streptomycin/amphotericin (Thermo Fisher) (RP10). CD4 T cells were plated in RP10 with plate-bound anti-CD3 (clone OKT3, eBioscience), anti-CD28/CD49d (BD Biosciences) and PHA-L (eBioscience) for 48h. NK cells were separately plated in RP10 with 300IU/ml recombinant human IL-2 (R&D) for 72h.

### In vitro HIV infection and NK co-culture assays

For all *in vitro* HIV infections, Q23-FL, a clone from early, subtype A infection (Poss and Overbaugh, 1999), was used. The Q23-FL virus was produced by transfecting a plasmid encoding a full-length, replication competent clone into 293T cells, harvesting supernatant after 48h and concentrating by ultracentrifugation. Viral stocks were titrated on TZM-bl cells as previously described (Strauss-Albee et al., 2015). Activated CD4 T cells were infected with Q23-FL at an MOI of 25 (based on titrations in TZM-bl cells), using Viromag magnetofection (OZ Biosciences). HIV-infected cells were used for co-cultures 24h post infection. NK cells and CD4 T cells were co-cultured at a 1:4 effector:target (E:T) ratio, for 4h, in the presence of brefeldin A (eBioscience), monensin (eBioscience), and anti-CD107a-APC (Biolegend).

### Mass cytometry

All antibodies were conjugated using MaxPar® X8 labeling kits (Fluidigm), except for those purchased directly from Fluidigm; details of all antibodies is given in Table S1. To maintain antibody stability and consistency in staining, all antibody panels were pre-mixed into separate surface and ICS cocktails (as indicated in Table S1), aliquoted and frozen at −80°C until use. Palladium (Pd102, Pd104, Pd106, Pd108) conjugated CD45 antibodies for barcoding were made as previously described (Mei et al., 2015).

At the end of co-culture, cells were stained for viability using 25 µM Cisplatin (Enzo) for 1min and quenched with FBS, and samples were barcoded using palladium-based CD45 barcodes as previously described (Mei et al., 2015). After barcoding, cells were washed thrice, and all samples from a set of barcodes were combined. Samples were stained with the surface antibody panel for 30min at 4°C, fixed with 2% paraformaldehyde (PFA, Electron Microscopy Sciences), permeabilized with Permeabilization Buffer (eBioscience), and stained with the intracellular staining (ICS) panel (made in Permeabilization Buffer) for 45min at 4°C. Cells were suspended overnight in iridium interchelator (DVS Sciences) in 2% PFA, and resuspended in 1x EQ Beads (Fluidigm) before acquisition on a Helios mass cytometer (Fluidigm).

### Data analysis

Bead normalization (https://github.com/nolanlab/bead-normalization) and debarcoding (using the R package *Premessa*) were performed on all files post-acquisition. All CyTOF data was visualized and gated using FlowJo v10.1 (Tree Star); gated NK cells (CD3^−^ CD56/CD16^+^), or functional^+^/functional^−^ cells were exported as fcs files from FlowJo and used in downstream analyses. CD11a was excluded from all downstream analyses due to poor staining. The data supporting this publication is available at ImmPort (https://www.immport.org) under study accession SDY1620.

The open source statistical software R (Team and Others, 2013) was used for analyses. For analyses using the generalized linear models as well as the clustering and the Uniform Manifold Approximation and Projection (UMAP) (McInnes et al., 2018), raw channel values were transformed using the inverse hyperbolic sine (asinh) function with a cofactor equal to 5 to account for heteroskedasticity. This transformation was not applied for calculating mean signal intensity values. To compare frequencies of functionally responding cells, as well as frequency of the gated population between HIV+ and HIV-groups, t-tests were used. To compare frequencies of responding cells between the gated population and bulk NK cells in each donor, paired t-tests were used. To identify markers that are predictors of the HIV+ or HIV-conditions, we used the R package *CytoGLMM* (Seiler et al., 2019) which uses a generalized linear model with bootstrap resampling. This model takes into account the distribution of each marker, and has a donor-specific variable to control for inter-individual variability. For the clustering analyses, the R package *CATALYST* was used (Nowicka et al., 2017; Weber et al., 2019). This package provides a clustering method which combines the *FlowSOM* algorithm (Van Gassen et al., 2015) which generates 100 high-resolution clusters, followed by the ConsensusClusterPlus metaclustering algorithm (Wilkerson and Hayes, 2010) which regroups these high-resolution clusters into metaclusters. Default parameters were used for clustering, and the number of metaclusters (10) was selected based on the delta area plot provided. To test for differential abundance of clusters between groups, the *diffcyt-DA-GLMM* method from the *diffcyt* package (Nowicka et al., 2017; Weber et al., 2019) was used; the donor IDs were specified as a random effect. The UMAP was run using the *scater* package (McCarthy et al., 2017), with default settings.

### RESULTS

### HIV+ donors have an altered NK cell repertoire even in the setting of ART suppression

To investigate the effect of HIV-1 infection on NK cell responses, we used CyTOF to profile NK cell receptor expression and functional activity in a cohort of 10 ART-suppressed, HIV+ donors (referred to as HIV+), and 10 healthy controls (referred to as HIV-). Patient demographics are given in Table 1. We first compared the expression of 28 NK cell receptors between the HIV+ and HIV-donors, in sorted, IL-2 activated NK cells. To look at overall NK receptor expression patterns between the two groups, we used a multidimensional scaling (MDS) plot to visualize all the NK cell samples (Fig 1A). NK cells from HIV- and HIV+ donors separated primarily on a diagonal axis; this separation was driven by multiple markers including CD2, NKp30 and NKp46 (as shown in the correlation circle). To further define the NK cell receptors whose expression is altered in HIV+ compared to HIV-individuals, we used a generalized linear model with bootstrap resampling to identify markers predictive of either the HIV+ or HIV-groups.

**Table 1:** Demographic information of HIV cohort

|  | HIV+ (n=10) | HIV- (n=10) |
| --- | --- | --- |
| Age in years \| Mean (SD) | 52.9 (9.7) | 54.1 (17.5) |
| Sex \| proportion male | 10/10 | 9/10 |
| Years since diagnosis \| Mean (SD) | 18.5 (8.6) | N/A |
| Years on ART \| Mean (SD) | 15.3 (8.1) | N/A |
| CD4 count in cells/mm<sup>3</sup> \| Mean (SD) | 705 (335) * | Not available |
| Nadir CD4 count in cells/mm<sup>3</sup> \| Mean (SD) | 249 (129) + | N/A |
| Type of ART (proportion) | NNRTI (6/10), PI (3/10), Integrase inhibitor (3/10) ^ | N/A |

\* CD4 counts were only available for 7/10 HIV+ individuals.
+ Nadir CD4 counts were only available for 6/10 HIV+ individuals.
^ NNRTI = non-nucleoside reverse transcriptase inhibitor, PI = protease inhibitor.

**Figure 1:**
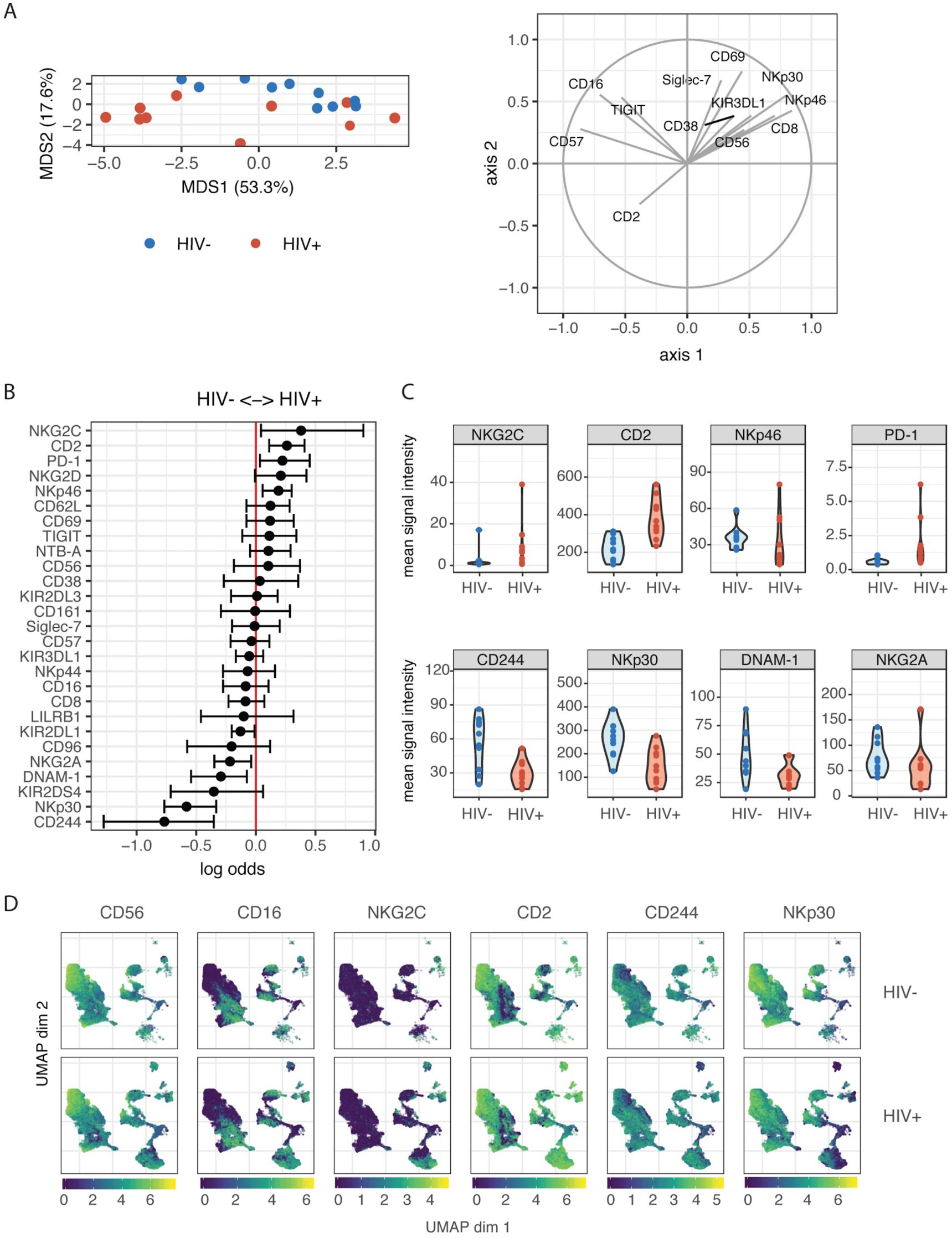
Profiling of NK cell repertoire using CyTOF in both healthy, HIV- and ART-suppressed, HIV+ donors demonstrates alterations in NK cell surface receptor expression. **(A)** Multidimensional scaling (MDS) plot showing separation of NK cells from HIV+ (n=10) and HIV-donors (n=10). Only markers whose contributions are greater than 0.25 in both MDS1 and MDS2 are displayed in the marker loadings. **(B)** A generalized linear model with bootstrap resampling was used to find receptors predictive of either HIV+ (right), or HIV-(left) donor NK cells. Log-odds are logarithm of ratios of the probability that a cell belongs to either group. For each marker, the 95% confidence interval is represented by the line surrounding the point estimate; a larger absolute log-odds value of the parameter indicates that the marker is a stronger predictor. **(C)** Mean signal intensity (MSI) of NKG2C, CD2, NKp46 and PD-1 (top 4 predictors of the HIV+ group; top), and CD244, NKp30, DNAM-1 and NKG2A (top 4 predictors of the HIV-group; bottom). **(D)** UMAP visualization of all NK cells from the HIV+ and HIV-groups, coloured by expression of CD56, CD16, NKG2C and CD2 (top 2 predictors of the HIV+ group), and CD244 and NKp30 (top 2 predictors of the HIV-group). Scales show asinh-transformed channel values.

Based on marker distribution, the model generates the log-odds that the expression of a given marker is predictive of either the HIV+ or HIV-group, together with the 95% confidence interval. In purified, IL-2 activated NK cells, the markers NKG2C, CD2, NKp46 and PD-1 were predictive of HIV+ (95% confidence interval does not contain the zero value) while CD244, NKp30, DNAM-1 and NKG2A were predictive of HIV-individuals (Fig 1B). To confirm these results, we also compared mean signal intensity (MSI) for the top 4 NK cell markers predictive of either the HIV-or HIV+ groups, and observed an increased trend of MSI for NKG2C, CD2, NKp46 and PD-1 in the HIV+ group, and a decreased trend in CD244, NKp30, DNAM-1 and NKG2A (Fig 1C).

To further visualize the changes in subsets of NK cells between HIV+ and HIV-groups, we used the Uniform Manifold Approximation and Projection (UMAP) to visualize purified NK cells from both groups (Fig 1D). To identify classic NK cell subsets, the expression of the markers CD56 and CD16 are shown, revealing the expected pattern in that the cells with highest expression of CD56 have low expression of CD16, identifying the canonical CD56^bright^CD16^−^ and CD56^dim^CD16^+^ NK cell subsets. In addition, the expression of the top predictors for each of the HIV-(CD244 and NKp30) and HIV+ groups (NKG2C and CD2) are shown. These plots reveal that greatest differences in both cellular distribution and NK marker expression between HIV-and HIV+ donors occurred on the right part of the UMAP plots. Thus, these data demonstrate that even in the setting of long-term virological suppression with ART, the NK cell repertoire following IL-2 treatment remains altered.

### NK cells from HIV+ individuals do not have an increased response upon in vitro restimulation with autologous HIV-infected cells

Contrary to their classic designation as innate immune cells, NK cells have more recently been shown to demonstrate antigen-specific memory to viral antigens. As such, we were interested in determining whether prior HIV-1 exposure (in the HIV+ individuals) would alter the magnitude of the NK cell response or the phenotypes of responding cells, when restimulated with autologous HIV-infected cells *in vitro*. We co-cultured NK cells from HIV- and HIV+ donors with autologous CD4 T cells infected *in vitro* with the HIV strain Q23-FL (Fig 2A); infection levels in CD4 T cells after co-culture were similar between HIV+ and HIV-donors (Figure S1). To identify NK cell responses, we looked at expression of functional markers on these cells by CyTOF. For each sample, we separately gated on NK cells that were positive for the expression of cytokines IFN-TNF-γ, MIP-1β (CCL4), TNF-α or the degranulation marker CD107a (Fig 2B), after 4 hours of co-culture. All gating was performed based on samples of NK cells in the absence of target cells; these samples had generally low levels of expression of all functional markers. We found that the majority of responding cells were polyfunctional: for example, using the generalized linear model, the top predictors of CD107a^+^ cells included MIP-1β, TNF-α and IFN-TNF-γ (Figure S2). As such, we used Boolean gating to identify functionally responding cells (positive for *any* of the functional markers above, hereafter named as functional^+^), or non-functionally responding cells (negative for *all* of the markers above, named as functional^−^).

**Figure 2:**
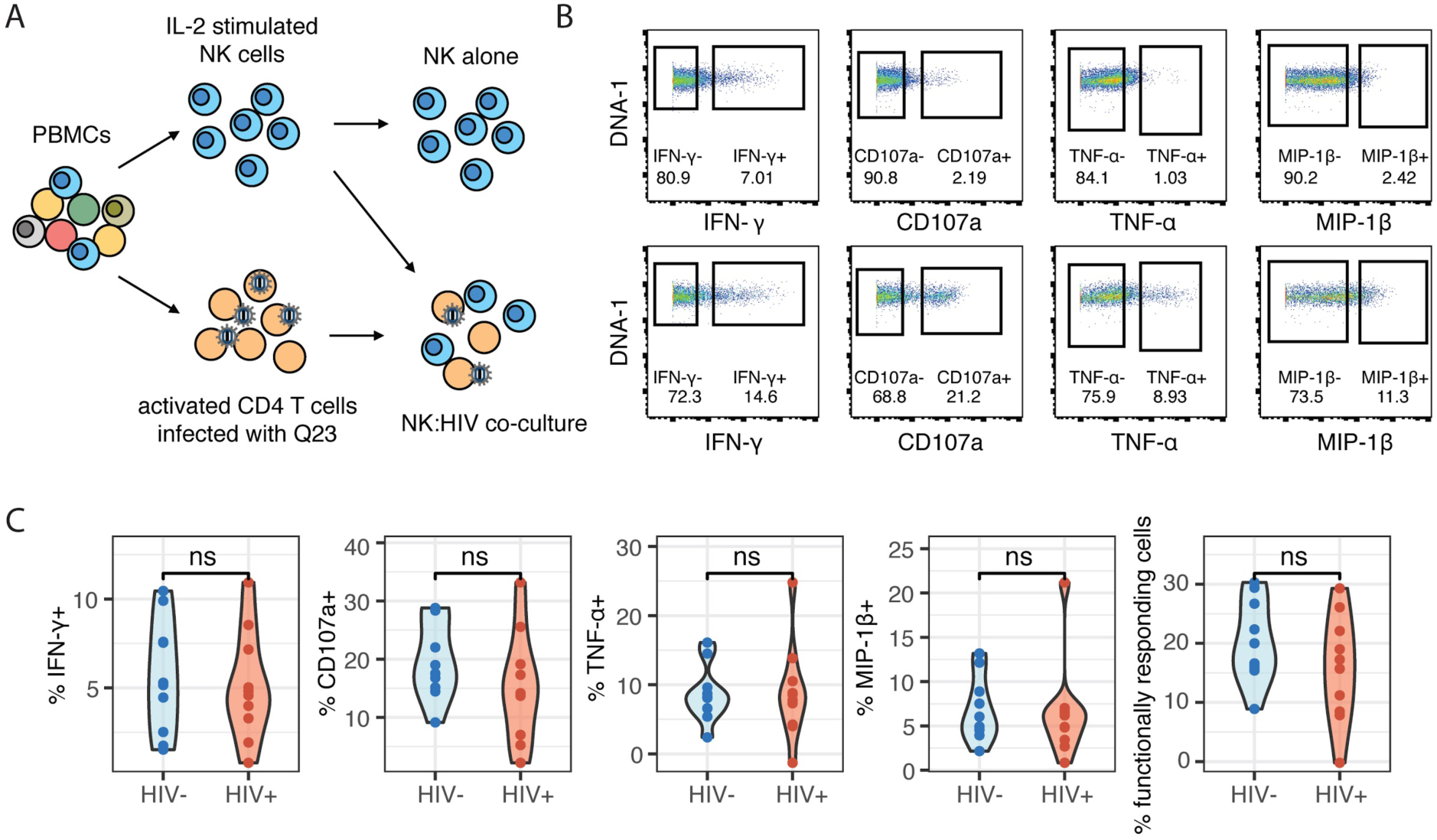
The magnitude of the HIV-specific response does not differ between HIV- and HIV+ donors. **(A)** Schematic of experimental set-up for NK co-culture assays. **(B)** Representative flow cytometry plots of CD107a, IFN-γ, MIP-1β and TNF-α production (by frequency of positive cells), after 4 hour co-culture without (top) or in the presence of HIV-infected autologous CD4^+^ T cells (bottom). **(C)** Summary data of CD107a, IFN-γ, MIP-1β and TNF-α production (by frequency of positive cells), after 4 hour co-culture with HIV-infected autologous CD4^+^ T cells, in NK cells from HIV-(n=10) and HIV+ (n=10) donors. ns = not significant, by t-test.

To identify differences in the magnitude of NK cell responses to HIV-infected cells between HIV- and HIV+ donors, we gated on cells that were positive for functional markers, and sought to identify differences in frequency of these cells between the two groups. We applied background subtraction (by subtracting the frequency of NK cells expressing each functional marker in the NK alone condition) to account for variations in baseline NK activity. In response to autologous HIV-infected cells *in vitro*, the frequency of NK cells expressing any of the functional markers (IFN-TNF-γ, MIP-1β, TNF-α, CD107a) individually, or combined (functional^+^), was not significantly different between the HIV- and HIV+ groups (Fig 2C).

To further understand whether the NK cells that were generating a functional response were phenotypically similar between the HIV+ and HIV-groups, we used the generalized linear model to identify predictors of responding (functional^+^) and non-responding cells (functional^−^). Multiple NK cell receptors were strong predictors of functional^+^ or functional^−^ cells, indicating a clearly distinct phenotype of cells that respond to HIV *in vitro*; these cells express higher levels of CD96, NKp30, TIGIT, and Siglec-7, and lower levels of CD62L, CD16, and NKp46 (Fig 3A). Notably, the top predictors of responding cells were very similar between the HIV- and the HIV+ group, including CD96, NKp30 and TIGIT. To identify a subset of responding cells, we included cells with high expression of positive predictors (predictors of functional^+^), and with negative expression of negative predictors (predictors of functional^−^), that had a log odds greater than 0.2 in both groups; the resultant phenotype was CD96^hi^ NKp30^hi^ TIGIT^hi^ CD16^−^ CD62L^−^ NKp46^−^. CD38, while being a strong predictor of functional^+^ cells in the HIV+ group, was excluded; it was not a predictor of responding cells in the HIV-group, and CD38 expression is increased in the context of HIV infection and remains high even after ART treatment (Kuri-Cervantes et al., 2014). We gated on each marker individually (Fig 3B), and then used Boolean gating to generate a gated population that included all the features of the phenotype. Gating on this subset significantly enriched for functional^+^ cells to a similar extent in both the HIV+ and HIV-groups (Fig 3C), suggesting that, despite the alterations in NK cell repertoire, the NK cell subsets responding to HIV *in vitro* were not altered in HIV+ individuals. We also compared the frequency of this subset among total NK cells, in both HIV+ and HIV-groups (Fig 3D), and noted that, while the frequency was low (between 0.1% and 3%), there was no significant difference between the two groups, indicating that HIV infection does not alter the frequency of this subset.

**Figure 3:**
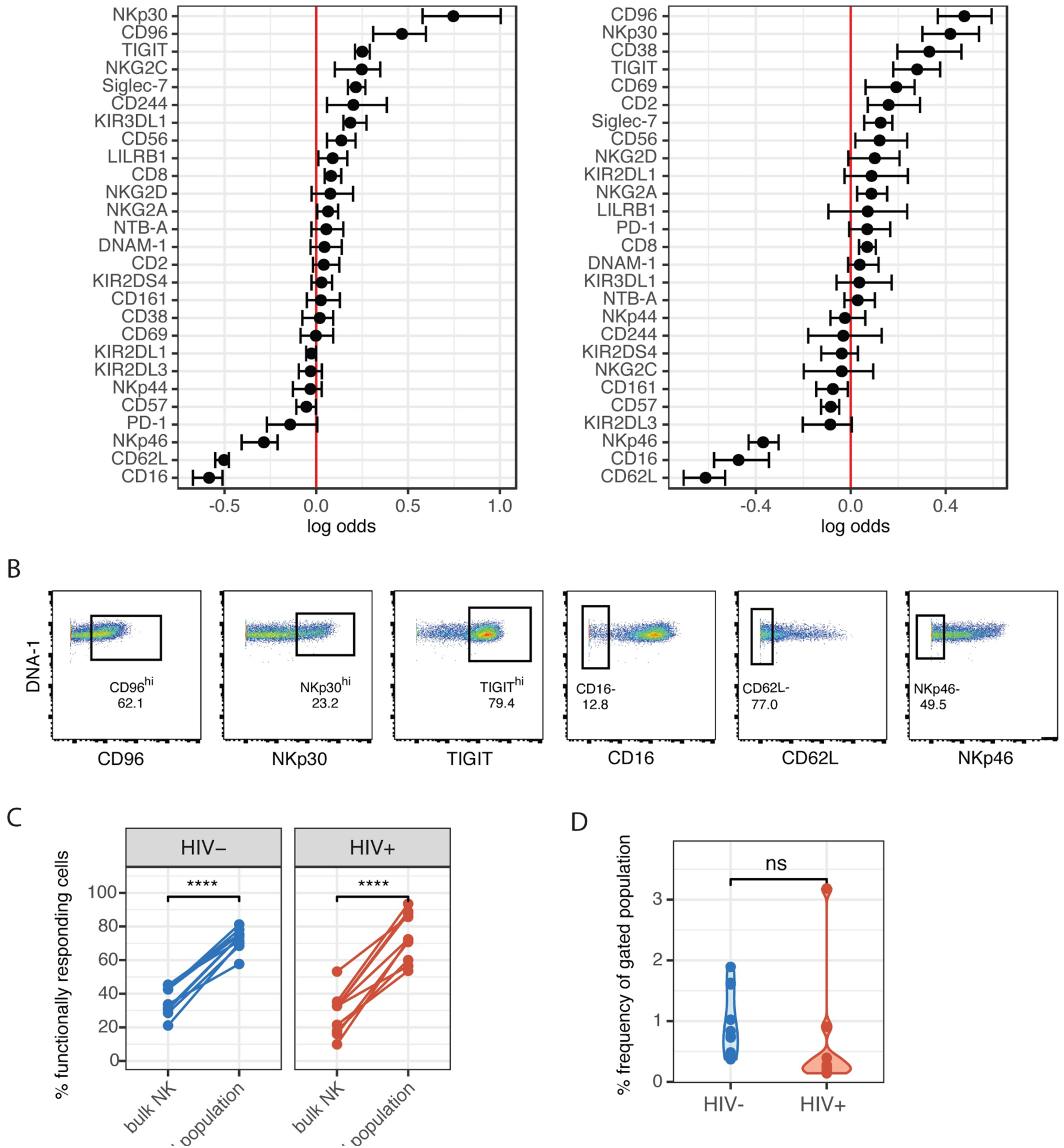
Phenotype of NK cells responding to HIV are similar between HIV- and HIV+ donors, and include many of the markers that are altered in ART-treated HIV infection. **(A)** A generalized linear model with bootstrap resampling was used to identify NK cell receptors predictive of either functionally responding (right), or non-responding (left) NK cells, in HIV-(n=10) and HIV+ (n=10) donors. Log-odds are logarithm of ratios of the probability that a cell belongs to either group. For each marker, the 95% confidence interval is represented by the line surrounding the point estimate; a larger absolute log-odds value of the parameter indicates that the marker is a stronger predictor. **(B)** Gating on markers that predict functionally responding cells (NKp30^hi^ CD96^hi^ TIGIT^hi^ CD62L^−^ CD16^−^ NKp46^−^), with a minimum log-odds threshold of 0.2 in both groups. **(C)** Comparison of functional activity in the gated population, compared to bulk NK cells. Summary data of percentage positive functionally responding (positive for any of CD107a, IFN-γ, MIP-1β and TNF-α) in HIV-(n=10) and HIV+ groups (n=10). **** = p < 0.001, by paired t-test. **(D)** Comparison of the frequency of the gated population, as a percentage of total NK cells, between the HIV-(n=10) and HIV+ (n=10) groups. ns = not significant, by t-test.

### NK cells clusters that are altered in HIV+ individuals are distinct from those that are functionally responsive to HIV in vitro

As the expression of many of the markers on responding cells were markers we found to be altered in the HIV+ group compared to HIV-(Fig 1), it was surprising that these changes did not seem to have an effect on the magnitude or quality of the NK cell response to HIV *in vitro*. To better understand the populations of cells that are altered in HIV+ individuals, compared to those that are involved in the functional response, we used UMAP to visualize all NK cells from co-cultures with HIV-infected cells *in vitro*, in both groups. We also performed unsupervised clustering and metaclustering, using the *FlowSOM* and *ConsensusClusterPlus* algorithms, to identify 10 metaclusters of NK cells (Fig 4A); this included clusters that were shared between HIV+ and HIV-groups, such as cluster 1, but also clusters that were distinct to NK cells from HIV+ individuals (cluster 3), or HIV-individuals (cluster 5). To identify responding NK cells, we overlaid marker expression of functional markers onto the UMAP visualization (Fig 4B). All responding NK cells (positive for functional markers) clustered in a similar area on the left part of the plots that was shared between HIV+ and HIV-donors; many of these cells are positive for multiple functional markers, confirming that most responding cells were polyfunctional (Figure S1). To identify which cluster these functionally responding cells belonged to, we looked at a heatmap of mean marker expression of all NK cell markers across all clusters (Fig 4C); this identified clusters 9 and 10 as the predominant clusters expressing functional markers, with cells in cluster 9 expressing high levels of CD107a, TNF-α and MIP-1β, and cells in cluster 10 also expressing high levels of IFN-TNF-γ. In line with our previous analysis of the phenotypes of functionally responding cells, cluster 9 expresses high levels of NKp30, while cluster 10 has relatively high levels of TIGIT expression and low levels of NKp46 expression; both clusters have low CD16 and CD62L expression.

**Figure 4:**
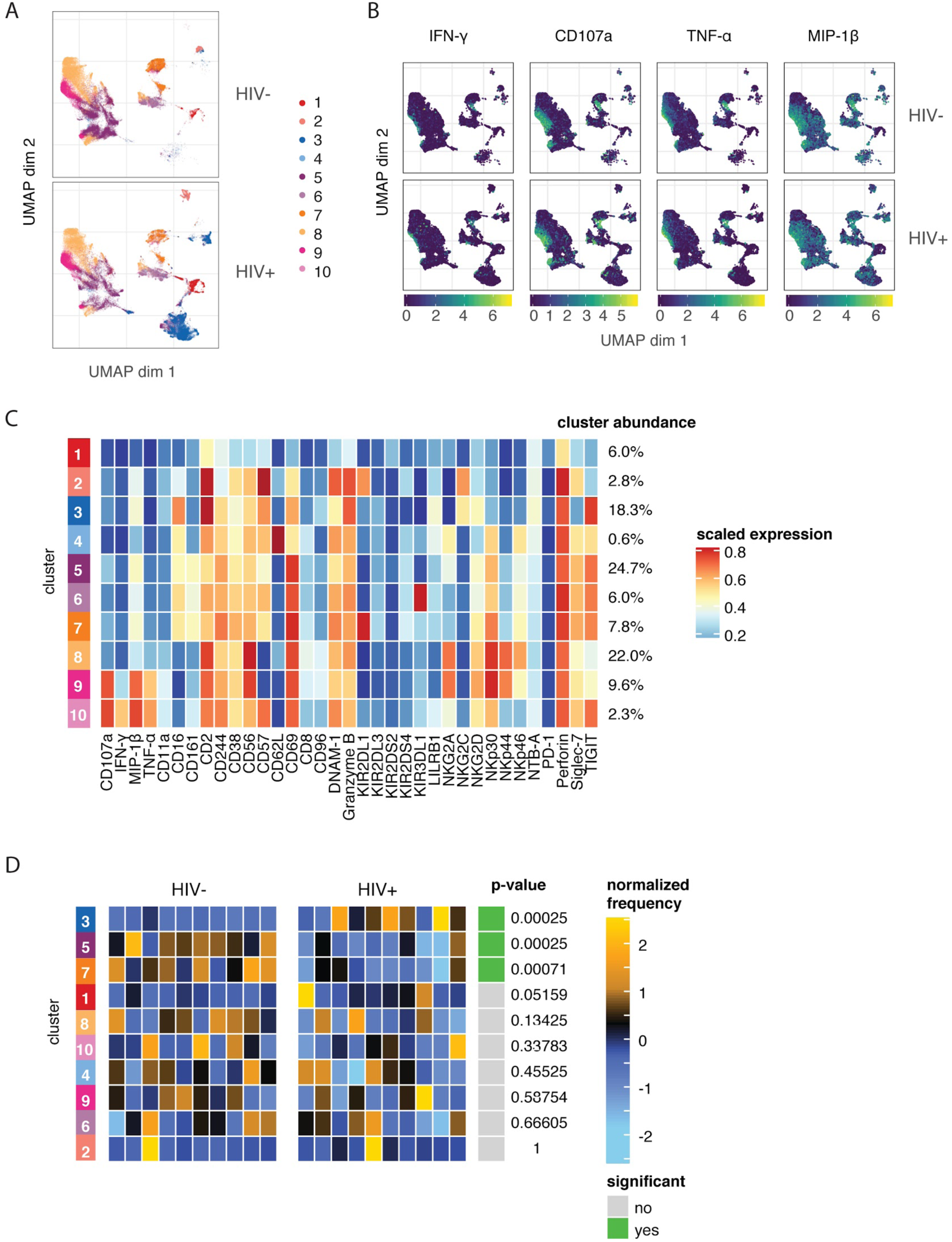
The predominant changes in the NK cell repertoire of HIV+ individuals occur in NK cell compartments that do not respond to HIV in in vitro restimulation. **(A)** UMAP visualization of all NK cells in co-culture with autologous HIV-infected cells in HIV-(n=10) and HIV+ (n=10) donors, coloured by metacluster identity generated by *ConsensusClusterPlus* metaclustering. **(B)** UMAP visualization of all NK cells from the HIV+ and HIV-groups, coloured by expression of functional markers CD107a, IFN-γ, MIP-1β and TNF-α. Scales show asinh-transformed channel values. **(C)** Heatmap of scaled mean expression of all markers profiled, for each cluster 1 to 10. The abundance of each cluster (% of total cells) is given on the right of the heatmap. Functional markers (CD107a, IFN-γ, MIP-1β and TNF-α) are on the left. **(D)** Heatmap of the relative abundance of each cluster between the HIV-(left) and HIV+ (right) groups. Each individual column represents a single donor. The heat represents the proportion of each metacluster in each donor, with yellow showing over-representation and blue showing under-representation. These proportions were first scaled with an arcsine-square-root transformation and then z-score normalized in each cluster. Clusters with a statistically significant (p<0.05) difference in abundance between HIV- and HIV+ groups are highlighted in green; adjusted p-values (FDR) are shown beside it.

To identify if these clusters were differentially abundant between the HIV+ and HIV-groups, we performed differential abundance tests using the *diffcyt* package, which identified 3 clusters with significantly different abundance between the HIV+ and HIV-groups - clusters 3, 5, and 7 (Fig 4D). Notably, neither cluster 9 or 10 were differentially abundant between the HIV+ and HIV-groups, indicating that the functionally responding subsets remain unaltered in abundance or phenotype (Fig 3) in the setting of ART-treated HIV infection. Hence, although HIV infection induces a reshaping of the NK cell repertoire *in vivo*, it does so in a way that does not alter HIV-specific responses, as the predominant changes occur in NK cell compartments that do not respond to HIV in *in vitro* restimulation.

## DISCUSSION

Chronic HIV-1 infection is known to alter NK cell phenotype and function. To better understand how these changes occur in the setting of virological control with ART, as well as how HIV-specific function is impacted in this setting, we used mass cytometry to profile differences in NK cell receptor expression repertoire in peripheral blood NK cells between ART-treated, HIV+ individuals and healthy HIV-controls. We observed differences in the IL-2 stimulated NK cell repertoire between HIV+ and HIV-individuals, although these differences did not impact the HIV-specific response to an *in vitro* restimulation with HIV-infected CD4 T cells. In addition, we identified a unique phenotype of cells that is functionally responsive against HIV; this NK cell subset is shared between both HIV+ and HIV-donors and is similarly responsive in both.

During HIV-1 infection, the NK cell repertoire undergoes significant changes, but even in the setting of virological suppression with ART, the NK repertoire remains altered compared to healthy controls in IL-2 activated NK cells. We found increased expression of NKG2C, CD2, NKp46 and PD-1, and decreased expression of CD244, NKp30, DNAM-1 and NKG2A; many of these markers have been previously described to be altered in HIV infection. The decreased expression of NKp30 and DNAM-1 has been reported for ART-treated HIV-infected patients compared to healthy donors (Bisio et al., 2013; Zhou et al., 2015); however, these studies also observed lower expression of NKp46 in HIV+ individuals, whereas we saw increased NKp46 expression in the HIV+ group. This discrepancy may be due to the IL-2 activation of NK cells in our study, which is known to increase NKp46 expression (Campos et al., 2015), and may do so to different extents in the HIV-and HIV+ groups. CD244 expression on NK cells is also known to decrease in HIV infection, although does recover over time after the initiation of ART (Ostrowski et al., 2005). PD-1 expression is also increased in HIV infection even with ART treatment, and these cells have limited proliferative capacity and may contribute to NK cell dysfunction (Norris et al., 2012). The increased expression of NKG2C and CD2 may reflect an increase in a subset of NK cells with a more mature, adaptive phenotype that is known to expand during HIV infection, and that persists even during ART treatment (Peppa et al., 2018). Indeed, we observed differential abundance of cluster 3 (Fig 4) between the HIV+ and HIV-groups; this cluster, which had greater abundance in donors of the HIV+ group, expressed high levels of CD2 and low levels of Siglec-7, and expressed both NKG2C and CD57, all known features of this subset.

We also identified other clusters of NK cells that are differentially expressed between HIV+ and HIV-groups (Fig 4). Clusters 5 and 7 have greater abundance in donors of the HIV-group; cells of these clusters have higher expression of CD244 and NKp30 in cluster 7, which we also identified in our GLM analyses (Fig 1B). These differentially abundant clusters may represent the loss of subsets of NK cells that occur after chronic HIV infection or ART treatment. The identification of these phenotypic features of NK cells can provide insight on how chronic, treated HIV infection shapes the activated NK cell repertoire, while the mechanisms and functional consequences of these alterations warrant further investigation.

We identified a unique NK cell subset that has higher functional activity in response to HIV-1-infected cells (Fig 3); this subset has the phenotype CD96^hi^ NKp30^hi^ TIGIT^hi^ CD16^−^ CD62L^−^ NKp46^−^, and its increased functional activity is present in both the HIV+ and HIV-groups. Individually, many of the markers represented in this subset have been previously implicated in immune-mediated control of HIV. For instance, in CD8 T cells, CD96 expression is positively associated with higher CD4 T cell counts in HIV-infected individuals, although CD96^+^ cells produce less perforin upon stimulation with phorbol myristate acetate/ionomycin (PMA/I), compared to CD96^−^ (Eriksson et al., 2012). In NK cells, CD96^+^ NK cells from peripheral blood have reduced TNF-α and IFN-TNF-γ production following PMA/I stimulation (Sun et al., 2019). These prior data make it difficult to ascertain whether CD96 contributes to HIV control, but our observation that CD96^hi^ cells have improved functional activity against HIV may reflect different mechanisms of activation between HIV infection and stimulation by PMA/I. In addition, NKp30 expression is induced upon IL-2 stimulation, and is correlated with IFN-TNF-γ production and inversely associated with the HIV reservoir, suggesting that cells that upregulate NKp30 upon IL-2 stimulation may have improved activity against HIV (Marras et al., 2017). This is particularly relevant as our study used IL-2 activated NK cells. TIGIT^+^ NK cells have also been previously implicated in HIV control -TIGIT expansion is markedly enhanced on NK cells in untreated HIV infection (Yin et al., 2018; Vendrame et al., 2020). We also recently demonstrated that TIGIT expression marks a population of NK cells with an adaptive phenotype with greater functional activity against HIV-infected cells as well as other stimuli (Vendrame et al., 2020).

In contrast, the lack of expression of CD16 and NKp46 on these functionally responding cells may reflect downregulation of these receptors after NK cell activation by HIV-infected cells. After stimulation, downregulation of both CD16 and NKp46 have been reported to occur predominantly in activated NK cells that produce IFN-TNF-γ and CD107a; this downregulation can occur even in the absence of specific signalling through CD16 or NKp46 (Grzywacz et al., 2007; Parsons et al., 2014). As such, these features of this subset may not necessarily reflect their involvement in the HIV-specific response, but instead mark activated NK cells. By using the combinatorial expression of all the markers we have identified, we were able to vastly enrich for cells that were responding against HIV -up to 90% of cells of this phenotype were functionally responding (Fig 3C). Although the frequency of this subset is low (0.1-3% of total NK cells, Fig 3D), and does not account for all responding cells, the high level of functional activity of these cells suggests that they are important in HIV-targeting activity.

While we did not find evidence of improved memory responses of peripheral blood NK cells in HIV+ individuals upon an *in vitro* re-stimulation with HIV-infected cells, this does not preclude the existence of these memory NK cells in other tissue locations, or in low frequency in the blood. In murine, humanized mice, and non-human primate studies of memory NK cells, memory NK cells were predominantly tissue-resident, particularly in the liver (Paust et al., 2010; Reeves et al., 2015; Nikzad et al., 2019). A low frequency of these hepatic phenotype NK cells can be found in peripheral blood (Nikzad et al., 2019) and would have been included in our analyses, but may have been at too low a frequency to detect within the bulk of the response to HIV. Indeed, Reeves et al. have previously found that, in non-human primates, memory responses to Gag vaccination by peripheral blood NK cells were much lower in magnitude than splenic or hepatic NK cells, but were still observable (Reeves et al., 2015). Even so, our observation that peripheral blood NK cells did not demonstrate HIV-specific memory responses may be due to the extremely low frequency of these cells, or differences between these memory NK cells in humans compared to mice or non-human primates or with different infections.

There are several limitations to our study. Due to sample availability, we profiled only NK cells in peripheral blood; tissue-resident NK cells may exhibit differences in both phenotype and the ability to generate memory responses, as discussed above. In addition, NK cells were not re-stimulated with the same HIV-1 strain as the primary *in vivo* infection, as we used a separate, *in vitro* infection with a different HIV strain. We used a subtype A strain for all *in vitro* HIV restimulations; however, as all our subjects were recruited in North America, where subtype B strains dominate (Bbosa et al., 2019), the mismatch in viral strain used for the secondary challenge may have contributed to the poor memory NK cell responses we observed. Even so, we have shown that infection with HIV-1 viruses across both subtype A and subtype B strains lead to similar patterns of alterations in expression of NK cell ligands on infected CD4 T cells (unpublished data), suggesting that strain-specific recognition is unlikely in NK cells, and the mismatch of strain subtype in primary infection and *in vitro* restimulation would not impair the detection of potential memory responses. Lastly, as our primary interest was in evaluating functional responses to HIV-infected cells, we profiled IL-2 activated NK cells, which may not entirely recapitulate NK cell phenotypes *ex vivo*. IL-2 induces changes in NK receptor expression, including upregulation of the natural cytotoxicity receptors (NCRs) NKp30 and NKp46; however, the expression of most other NK receptors remains unchanged (Vendrame et al., 2017). The differences in cytokine-induced upregulation of these NCRs between the HIV+ and HIV-groups can additionally be informative, as upregulation of NCRs have been implicated in control of HIV (Marras et al., 2017).

In summary, our data demonstrate that phenotypic alterations in peripheral blood NK cells that occur in individuals with ART-treated HIV-1 infection do not result in improved NK-mediated targeting of HIV. These phenotypic changes instead occur in distinct cellular subsets that are not involved in the functional response to HIV. Further work is required to understand whether other tissue-resident NK cells may exhibit differences in phenotypic alterations and functional responses in the course of treated HIV infection.

## ACKNOWLEDGEMENTS

We would like to thank Thanmayi Ranganath for assistance with running mass cytometry samples, and the Human Immune Monitoring Core (HIMC) at Stanford University for use of their Helios machine.

## AUTHOR CONTRIBUTIONS

NZ and CB designed experiments. NZ conducted experiments. NZ and AF analyzed the data with statistical analysis input from SH. PG contributed samples to the study. NZ and CB wrote the manuscript; all authors contributed revisions.

