## Supplemental data for "Treated HIV Infection Alters Phenotype But Not HIV-specific Function of Peripheral Blood Natural Killer Cells"

**SUPPLEMENTARY INFORMATION**

*Table S1: Antibodies used for mass cytometry*

| Isotope | NK Marker | Source | Clone | Panel |
|---|---|---|---|---|
| 89Y | CD57 | Biolegend | HCD57 | Surface |
| 112Cd/Qdot | CD19 | Invitrogen | SJ25-C1 | Surface |
| 115In | CD3 | Biolegend | UCHT1 | Surface |
| 141Pr | Granzyme B | Invitrogen | GB11 | ICS |
| 142Nd | MIP-1β | BD Biosciences | D21-1352 | ICS |
| 143Nd | NKG2C | R&D Systems | MAB1381 | Surface |
| 144Nd | CD161 (KLRB1) | BD Biosciences | DX12 | Surface |
| 145Nd | CD38 | Biolegend | HIT2 | Surface |
| 146Nd | CD8 | Biolegend | SK1 | Surface |
| 147Sm | CD107a (LAMP1) | Biolegend anti-APC | APC003 | ICS |
| 148Nd | LFA-1 (CD11a/CD18) | Biolegend | M24 | Surface |
| 149Sm | CD2 (LFA-2, LFA-3) | Biolegend | RPA-2.10 | Surface |
| 150Nd | HIV p24 core antigen | abcam | 39/5.4A | ICS |
| 151Eu | Siglec-7 | Biolegend | S7.7 | Surface |
| 152Sm | Perforin | abcam | B-D48 | ICS |
| 153Eu | KIR2DS4 (CD158i) | R&D Systems | 179315 | Surface |
| 154Sm | LILRB1 (ILT-2/CD85j) | R&D Systems | 292319 | Surface |
| 155Gd | NKp46 (CD335) | Biolegend | 9E2 | Surface |
| 156Gd | NKG2D | Biolegend | 1D11 | Surface |
| 157Gd | TIGIT | R&D Systems | 741182 | Surface |
| 158Gd | CD244 (2B4) | Biolegend | C1.7 | Surface |
| 159Tb | CD226 (DNAM-1) | BD Biosciences | DX11 | Surface |
| 160Gd | IFN-γ | BD Biosciences | B27 | ICS |
| 161Dy | NKp30 (CD337) | Biolegend | P30.15 | Surface |
| 162Dy | TNF-α | eBioscience | MAB11 | ICS |
| 163Dy | KIR3DL1 | BD Biosciences | DX9 | Surface |
| 164Dy | NKp44 | Biolegend | P44.8 | Surface |
| 165Ho | CD96 (TACTILE) | Biolegend | NK92.39 | Surface |
| 166Er | KIR2DL1 | R&D Systems | 143211 | Surface |
| 168Er | CD62L | Biolegend | DREG-56 | Surface |
| 169Tm | NKG2A | Fluidigm | Z199 | Surface |
| 170Er | KIR2DS2 | Abcam | Polyclonal | Surface |
| 171Yb | PD1 (CD279) | Biolegend | EH12.2H7 | Surface |
| 172Tb | NTB-A | Biolegend | NT-7 | Surface |
| 174Yb | CD56 | BD Pharmingen | NCAM16.2 | Surface |
| 175Lu | KIR2DL3 | R&D Systems | 180701 | Surface |
| 176Yb | CD69 | Biolegend | FN50 | Surface |
| 209Bi | CD16 | Fluidigm | 3G8 | Surface |

*Figure S1: Levels of HIV infection on CD4 T cells is similar between HIV+ and HIV- groups after co-culture with NK cells.*

Summary data of infection levels of CD4 T cells (by frequency of cells staining for HIV p24), after 4 hour co-culture with HIV-infected autologous $CD4^{+}$ T cells, in NK cells from HIV- (n=10) and HIV+ (n=10) donors. ns = not significant, by t-test.

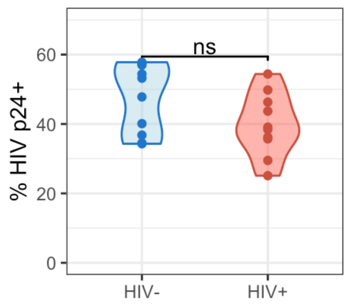

*Figure S2: Functional markers are co-expressed on many responding NK cells.*
A generalized linear model with bootstrap resampling was used to identify NK cell receptors predictive of either CD107a$^{+}$ (right), or CD107a$^{-}$ (left) NK cells, in HIV- (n=10) and HIV+ (n=10) donors. Log-odds are logarithm of ratios of the probability that a cell belongs to either group. For each marker, the 95% confidence interval is represented by the line surrounding the point estimate; a larger absolute log-odds value of the parameter indicates that the marker is a stronger predictor.

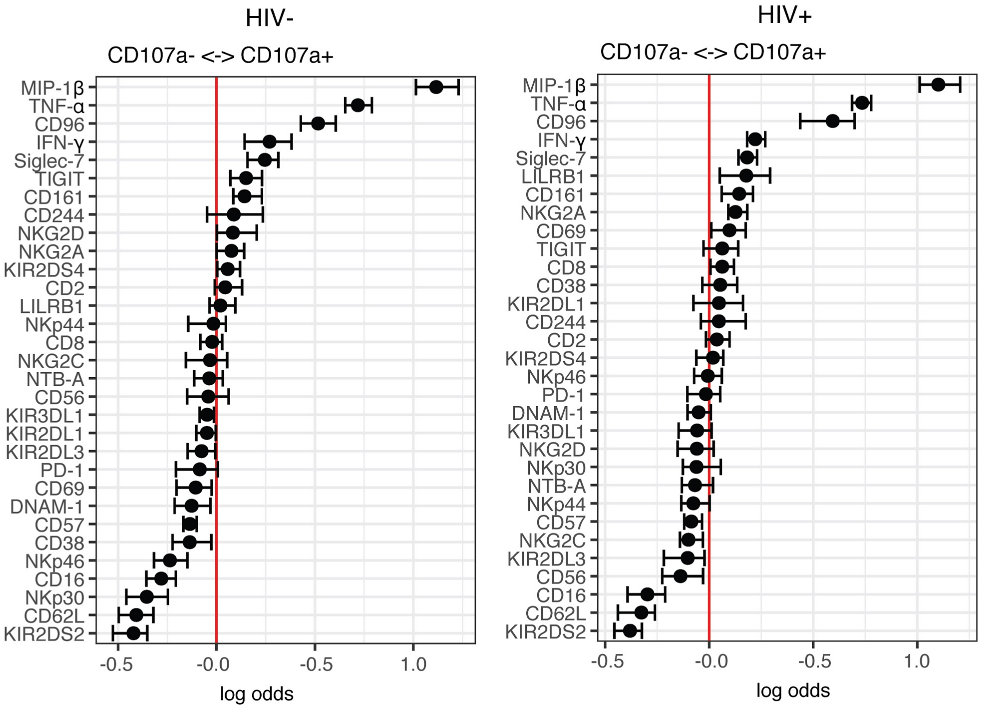
